# Stability of eye movement-related eardrum oscillations to acoustic and gravitational/postural manipulations

**DOI:** 10.64898/2026.04.16.718961

**Authors:** Nancy Sotero Silva, Christoph Kayser

**Affiliations:** Cognitive Neuroscience, Faculty of Biology, Bielefeld University, Bielefeld, Nordrhein-Westfalen, Germany

**Keywords:** eye movements, saccades, tympanic membrane, otoacoustic emissions, sensory-motor integration

## Abstract

Recent studies describe Eye Movement-Related Eardrum Oscillations (EMREOs), low-frequency signals recorded in the ear canal that arise from the tympanic membrane and are triggered by saccadic eye movements. Because EMREOs are thought to arise from motor elements in the peripheral auditory system, we examined how two known modulators of these elements affect the EMREO time course. First, the activity of outer hair cells (OHC) can be suppressed by the medial olivocochlear reflex (MOCR). If OHCs contribute to the generation of EMREOs, activation of this reflex should reduce EMREO amplitude. To test this, we compared EMREO amplitudes elicited by saccades performed in silence and in the presence of contralateral noise. Second, postural (i.e. vestibular and proprioceptive) cues linked to head orientation may influence EMREOs via oculomotor control circuits that possibly modulate middle ear muscles. To test this, we recorded EMREOs while participants made saccades with their head upright (0° azimuth) and with their head tilted 30° in either direction. Across both experiments our data reveal no clear modulation of the EMREO time course by these experimental manipulations. Together with other recent studies these findings advocate for a stability of the EMREO time course towards multiple experimental modulations and fuel speculations that the signal may serve as a temporal reference frame rather than a spatial coordinate transform function when combining signals across the senses.

## 1. Introduction

Eye movement-related eardrum oscillations (EMREOs) are low-frequency signals recorded from the tympanic membrane following saccadic eye movements [1]. Recent studies describe that EMREOs last for up to 100 ms after saccade onset and exhibit deflections of opposing sign and distinct timing for saccades in different directions along the horizontal and vertical dimensions [2–5]. In particular, the EMREO induced by horizontal saccades is a reliable signal that carries information about saccade direction and possibly also amplitude [4,5]. However, the precise origin of the EMREO and the neural pathways and structures giving rise to this phenomenon remain unclear. The EMREOs are likely generated by motor components in the middle ear and the cochlea. Particularly the middle ear muscles (MEMs) and the outer hair cells (OHCs) have been indicated as key candidates [1,5,6]. The MEMs and OHCs play a role in actively adjusting sound transduction in response to sudden loud sounds. However, the underlying reflexes and feedback pathways also receive information about other motor-related signals. Self-vocalization, eye and head movements associated with shifts in attention, or body posture [7] are visible as contextual or cross-modal influences on auditory reflexes [8–10]. These feedback signals entering the pathways to the OHCs and MEMs likely induce the EMREO in a saccade-specific manner. However, they may in principle also shape the EMREO by additional sensory or body-related information that is not directly tied to the execution of a saccade [11–13]. We here probed for such non-saccade contextual effects on saccade-induced EMREOs in the human ear: first, we tested for an influence of acoustic context by adding a contralateral suppression noise, and second, we tested for an influence of non-acoustic context by tilting the head.

The OHCs help protect the cochlea from external noise, improve speech-in-noise perception, and facilitate spatial hearing [14–16]. Previous studies suggest that the OHCs may also receive information about oculomotor behavior that can possibly contribute to the generation of the EMREO [5,17]. These signals involve descending information from the superior colliculus via the superior olivary complex, as determined by animal studies [18–23]. However, signals induced by OHC activity, such as otoacoustic emissions, are generally much smaller than typical EMREOs [1]. Still, one could expect that experimental manipulations interfering with normal OHCs function may also affect the EMREO, in particular if a contribution to the EMREO signal arises from OHCs. One way by which one can interfere with OHC function is the medial olivocochlear reflex (MOCR)[24,25]. This can be triggered by a suppression noise, which when presented to one ear induces a hyperpolarization of the OHCs in the opposing ear [15,26,27]. The effect of this suppression can be seen in the reduction of transient evoked otoacoustic emissions (TEOAEs)[2,16,28]. These are transient sounds generated by the OHCs in response to brief acoustic stimuli, which can be routinely recorded using in-ear microphones and are used in clinical screening [29]. We here tested whether inducing the MOCR, as confirmed by a suppression of contralateral TEOAEs, alters the time course of the EMREO induced by horizontal saccades in the opposing ear. Our assumption was that if OHCs contribute to generating the EMREO, any process that impacts OHCs activity (e.g., the inhibitory effect posed by MOCR) should also alter the EMREO time course in a reliable manner.

The MEMs, i.e., the stapedius and tensor tympani, are connected to the tympanic-ossicular system and adjust sound transmission in loud environments via the acoustic reflex [9,30–32]. Importantly, the pathways mediating the acoustic reflex involve structures that receive multisensory and motor-related signals, whose anatomical origin remains partly unknown [7,33–35]. Again, these pathways offer a possible route for oculomotor signals to shape middle ear function and may contribute to the EMREO. In fact, non-oculomotor signals have been shown to activate the MEMs, thus reducing the response from the auditory system to body sounds (self-voice, heartbeats, joints creaking)[36]. Since oculomotor control in everyday situations requires the combination of eye, head and body-related postural spatial information [37–40], one could assume that the pathways providing motor-related information to the MEMs also carry information related to the orientation of the head in space to the middle ear. For example, the extraocular muscles receive feedback from the vestibular system that is taken into account when coordinating eyes and head during gaze control. Moving the head in relation to the body triggers vestibular reflexes and activates the various eye muscles in a differential manner [38,41,42]. Frontal-plane head tilts provoke excitatory stimulation in the semicircular canals of the ipsilateral ear and inhibitory stimulation in the contralateral ear. The signal transmitted by the vestibular nuclei leads to activation of the oculomotor and trochlear nerves, generating contraction of the superior rectus and oblique muscles in the ipsilateral eye, and inferior rectus and oblique muscles in the contralateral eye, rotating the eyes opposite to the head orientation [43]. In addition, changes in body posture have been shown to induce subtle changes in hearing thresholds and otoacoustic emissions, induced by changes in cochlear fluid pressure and by triggering contractions of the MEMs [10,44]. We tested for the presence of such head-posture-related vestibular and proprioceptive information in the pathways giving rise to the EMREO. For this we compared EMREOs between conditions that differ in their vestibular context by having participants tilt their head to the left or right.

Collectively, the present study tested whether two distinct contextual factors, the presence of contralateral suppression noise and changes in head-tilt, shape the EMREO time course. Our primary goal was to determine whether these factors have an influence on the EMREO time course in humans, information that helps constrain our understanding of its putative function and the neural pathways giving rise to this phenomenon. Our goal was not to determine directly whether or how strongly the OHC or MEMs in isolation contribute to the EMREO, and quantifying their individual contribution would require experiments with additional and more invasive manipulations. For our study we recorded EMREO signals in human participants in two experiments that required them to perform visually guided saccades (Fig 1). In one experiment, participants performed horizontal saccades while we measured EMREOs in the presence and absence of a contralateral suppression noise that successfully reduced TEOAEs in the ipsilateral ear. In the second experiment, the same participants performed saccades along horizontal, vertical and diagonal directions with their head straight, or tilted to the left or right. Overall, our results demonstrate a stability of the EMREO properties and time course across the tested conditions. This suggests that the EMREO constitutes a phenomenon that is tied to the execution of a saccade but is otherwise robust to various contextual manipulations.

**Fig 1.**
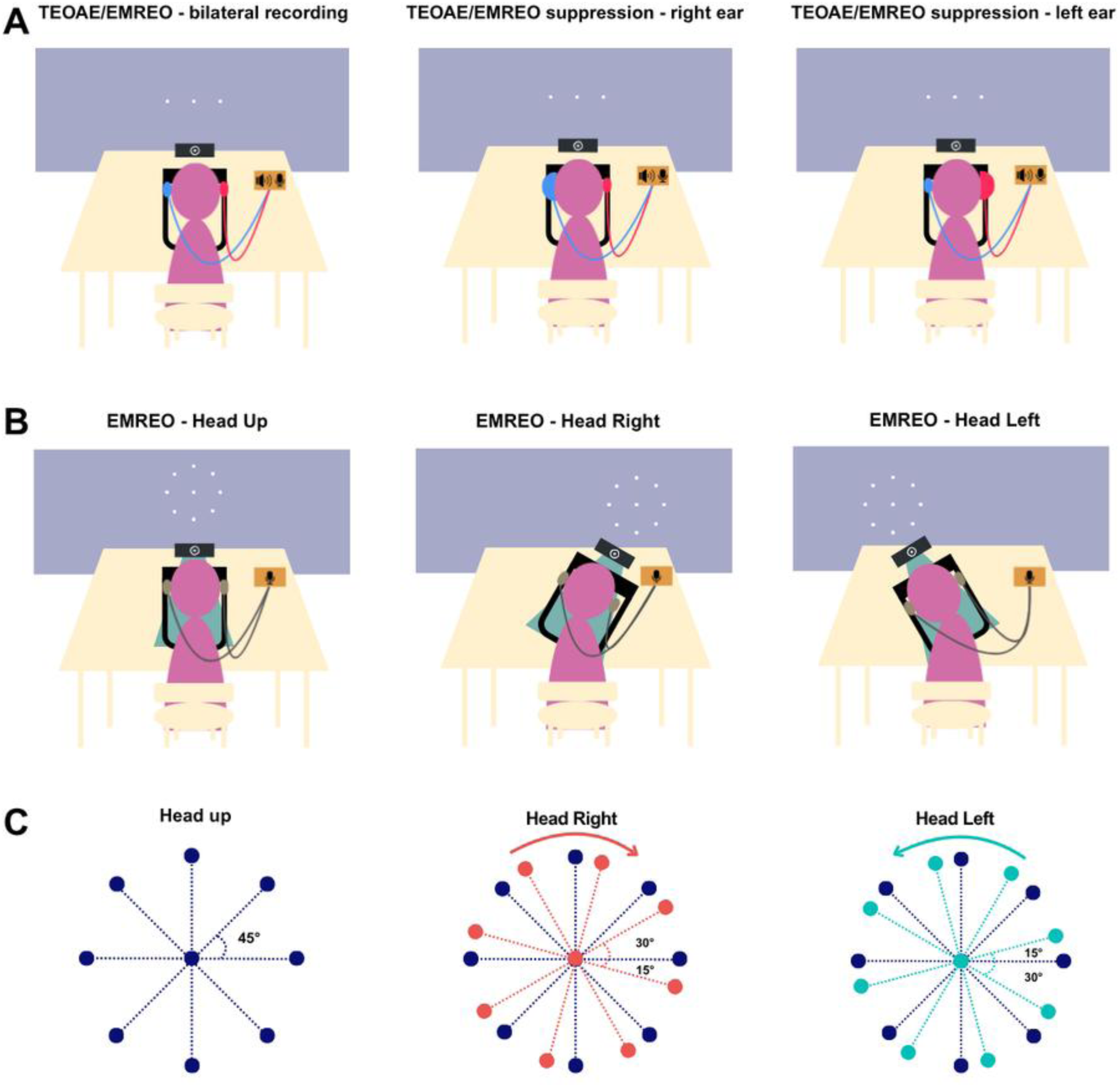
Schematic representation of the experimental paradigms. (A) The setup used in experiment 1 for recording TEOAEs and EMREOs for horizontal saccades. Participants sat in front of a screen onto which saccade targets were projected while we recorded eye movements and in-ear signals. From left to right: in different blocks we either recorded bilateral TEOAEs or EMREOs, or recorded these signals in one ear (smaller ear probe) while the other ear was suppressed using noise (larger headphone). (B) The setup used in experiment 2 for recording EMREOs during head-tilts for saccades along cardinal and oblique axes. From left to right: we recorded EMREOs while participants had their head upright or tilted to the right or left. For this, the chinrest and the eye tracker were mounted on a tiltable platform. (C) Schematic of saccade target positions in experiment 2 for the different conditions. Saccade targets were presented along the horizontal, vertical and oblique dimensions on the screen (dark blue). In the Head Right and Head Left conditions participants’ heads were rotated by 30° to the right and left respectively, inducing a relative rotation of head and screen coordinates. Arrows indicate the direction of the head tilt.

## 2. Materials and methods

### 2.1. Participants

A total of 47 participants of both sexes were recruited through a dedicated online platform among the university students and community members (31 female, 16 male, mean ± SD age = 30,19 ± 9,75). The sample size was planned based on previous studies studying the properties of EMREOS [2,3,31,45]. The study was approved by the Bielefeld University Ethics Review Board (#2024-037). All participants provided written informed consent, reported no history of hearing or neurological disorders and were compensated with 15 Euros per hour. Participants first underwent hearing screening, consisting of a visual inspection of the ear canal, Pure Tone Audiometry (PTA), tympanometry and testing of acoustic reflexes, as described previously in detail [2]. Participants with hearing loss (PTA average of 500, 1000, 2000 and 4000Hz above 20 dB) or middle ear impairments (as ear canal obstructions, membrane curves type B or C) in at least one ear were excluded from the study. We also tested participants’ visual acuity, and accepted participants with uncorrected impairment of < ± 2 diopters. Participants were given the option to perform the experiment with or without glasses or contact lenses. From the 47 recruited participants, four did not proceed to the actual data collection stage: two subjects due to high hearing thresholds, and two as the eye tracking system did not reliable track their eyes. Hence, we collected EMREO data from 43 participants.

### 2.2. Experimental setup

Participants were recruited and the experiments were conducted between April and July 2025. The experiments were carried out in a soundproofed, dark and electrically shielded booth. Participants were seated 100 cm away from a sound-permeable screen (Screen International Modigliani, 2 × 1 m) with their head on a chinrest aligned to the center of the screen. In-ear microphone recordings were performed using two Etymotic ER-10C (Etymotic Research) systems, via probes that served both for sound presentation (to induce TEOAEs) and in-ear recordings (TEOAEs and EMREOs). While recording the EMREO from a given year, the respective acoustic input to that ear was disconnected from the audio-amplifier to avoid any potential contamination of the recorded EMREO signal. Acoustic stimuli for TEOAEs (clicks) and noise for contralateral suppression were created using a Sound Blaster X3 Soundcard (Creative Technology Ltd., sampling rate 48’000 Hz) and amplified with a HeadAMP 4 (ARTcessories) amplifier. Click stimuli were presented via the ER-10C probes and the noise was delivered via a unilateral headphone (Radio Ear DD45C headphones, model HB3045). The recorded microphone signals were pre-amplified using the ER-10C DPOAE amplifiers (Etymotic Research) with gain set to +20 dB. Signals were digitized through an ActiveTwo AD-box (BioSemi) at a sampling rate of 16’388 Hz. The intensity of the acoustic stimuli was measured with a Hand-held Sound Level Meter (Model 2250, Bruel & Kjær, Denmark) using Instantaneous Peak Sound Level parameters (LCpeak, 1s). Visual stimuli were projected (Acer Predator Z650, Acer; 60 Hz refresh rate) onto the screen. Stimulus presentation was controlled using the Psychophysics toolbox [46] for MATLAB [47], which was synchronized to the recording system using transistor-transistor logic (TTL) pulses.

Eye movements were recorded at 1000 Hz from the left eye using a desktop mounted infrared-based EyeLink 1000 Plus Eye Tracker [48] running Host Software vs 5.15. Eye tracking calibration and validation was performed at the beginning of each block using a 5-point grid. The parameters for saccade detection in the EyeLink system were a velocity threshold of 30°/s and an acceleration threshold of 8000°/s² (“cognitive” setting). Eye position was recorded as horizontal and vertical position coordinates in degrees. In all experiments the eye tracker camera was aligned with the chinrest and participants’ head. Fig 1 presents a schematic representation of the experimental setup and paradigms for Experiment 1 (Fig 1A) and Experiment 2 (Fig 1B, C).

We performed control measurements for the in-ear microphones to ensure system calibration prior to each session. These recordings included an empty-room measurement consisting of 15 seconds of silence and the presentation of 20 tone stimuli (10 stimuli at 100 Hz, 10 stimuli at 200 Hz, duration 400 ms) presented from the speakers behind the screen. For this empty-room measurement, the in-ear probes were attached to the chinrest to simulate their typical (in-ear) position related to participants’ heads. The same measurement was repeated with the in-ear probes inserted into the participants’ ear canals. The baseline noise spectrum and the signal amplitudes in response to these tones were used to adjust the ear-probe insertion and to calibrate the signal levels between ears, as in our previous studies [2].

### 2.3. Measurement and suppression of TEOAEs

To induce and measure TEOAEs we followed established procedures [49]. The paradigm presented a series of 600 binaural clicks (rectangular monopolar signals of 0.1 ms duration, delivered at 83 ± 2 dB) presented with inter-click intervals of 380 to 420 ms (uniform, random). Every fourth click had an inverted polarity and three times the amplitude of the standard clicks. Participants were instructed to sit and listen as quietly as possible and to not move their head.

To induce the suppression of TEOAEs, we presented the same clicks to one ear while presenting a suppression noise to the other ear. The suppression noise consisted of a white noise (sampled at 48’000 Hz, delivered at 71 ± 2 dB) presented continuously for the presentation of 8 clicks followed by a silent interval of the same duration. The intensities for the click and noise were chosen based on established procedures [50–52]. For each participant TEOAEs were first measured without suppression, then with unilateral suppression. For the latter, during the second and third blocks we presented the clicks ipsilaterally and the suppression noise contralaterally with the in-ear probes. The order of which ear was suppressed first was counterbalanced across participants.

### 2.4. Measurement and suppression of EMREOs

We collected EMREOs during visually-guided horizontal saccades, similar to our previous studies [2,3]. As in the TEOAEs paradigm, we first measured baseline EMREOs, then EMREOs in one ear while the other ear was presented with noise to induce TEOAE suppression (Experiment 1, Fig 1A). The visual saccade task comprised horizontal saccades either from a central fixation position to a left (-12°) or right (12°) target, or back from these eccentric targets to the central position. In each trial, the fixation dot (0.2° radius, white on the gray background) was presented steadily for 650 ms at the respective start location before it jumped either to one of the two eccentric targets or back to the central position. The dot remained on the screen continuously. Each eccentric position was presented 40 times, the central position 80 times, resulting in 160 saccades per block. Intertrial intervals lasted between 750 and 1050 ms (uniform). Participants were instructed to fixate their gaze on the dot and to follow this as quickly as possible. For data analysis, we separated trials by saccade direction (left, right), for which we obtained 80 trials each. During the second and third blocks one ear was suppressed using noise, while EMREOs were recorded from the other ear. For this, we substituted one of the in-ear probes with the unilateral headphone over which we presented the same type of noise and with the same calibrated amplitude as used to suppress the TEOAEs (Fig 1A).

### 2.5. Measurements of EMREOs during head tilts

To test the influence of head-tilt on EMREOs (Experiment 2), we collected EMREOs during visually-guided saccades along horizontal, vertical and diagonal dimensions. The paradigm was similar as in Experiment 1, but here the fixation target jumped from the central position to one of eight eccentric positions spaced at 45° from each other (at 8° distance from the center). In alternating trials, the fixation dot either jumped to an eccentric target (in a pseudo-random sequence) or back to the center. Each eccentric position was presented 15 times per block, the central position 120 times, resulting in a total of 240 saccades per block, leading to 720 trials in total.

The main rationale of this experiment was to test whether tilting the head would influence the EMREO signal. For this experiment we mounted the chinrest and the eye tracker onto a platform that could be tilted and secured in three different positions: in the first block the platform was kept at 0° (head upright). For the second and third blocks, the platform was tilted 30° to the right or the left (order counterbalanced across participants). Participants were instructed to keep the body as straight as possible and only tilt their neck to adjust the head position. For the head-tilted conditions, the stimulus projection was adjusted on the screen (lowered and moved to the side of the tilt) to ensure that the center fixation was at the same level to participants’ eyes in each block, but the targets’ locations were kept upright for all conditions (see Fig 1B). Because in the tilted blocks the axis of the head and the calibration grid on the screen were rotated relative to each other, we transformed all eye positions into head-centered coordinates for data analysis. Note that although the degree of head tilt (30°) was not the same as the spacing of saccade directions (45°), this difference does not constrain the interpretation of the results (see below in the Results session, also Fig 1C). The head tilt implements an experimental condition that misaligns world and head axes and induces differences in vestibular and body-related postural signals.

The processed TEOAE and EMREO data included in the analysis is available publicly in the Bielefeld University Publications repository (https://doi.org/10.4119/unibi/3018705). The MATLAB codes used in the analysis are available as a Bielefeld University GitLab repository (https://gitlab.ub.uni-bielefeld.de/nsoterosilva/stability_emreos_acoustic_postural/).

### 2.6. Analysis of TEOAEs

The microphone signals were converted to mPa sound pressure based on the amplifier settings and the specifications of the in-ear probes provided by the manufacturer. The signals were high-pass filtered at 600 Hz and epoched around the individual clicks in windows of -25 to 30 ms. Epochs whose standard deviation exceeded 1.7 the standard deviation of the collective data across all epochs were discarded. To obtain an estimate of the noise level in the signal, we extracted the same number of epochs from randomly selected periods in between clicks (termed “baseline epochs”), which were devoid of any acoustic stimulus or response of the auditory system (ensuring a distance of > 50 ms from each click). We then analyzed the TEOAE and baseline epochs in the frequency domain. For this we calculated average signal power between 600 to 5000 Hz in steps of 66 Hz across epochs (FFT with Hanning taper, zero-padded to the next power of two). For each participant we then computed the difference in signal power between clicks and baseline, to obtain an estimate of the effective TEOAE in the frequency domain. This TEOAE spectrum was obtained separately for each block (considering only the non-noise ears). For each ear, we then compared the group-level spectra between blocks without and with suppression (see below in “Analysis of saccade characteristics”). To summarize the TEOAE signal amplitude, we integrated the difference between signal and baseline spectra between 576 Hz and 1856 Hz for each channel and participant.

### 2.7. Analysis of saccade characteristics

To ensure that the saccades driving the EMREOs in the different experimental conditions are comparable in their characteristics (silence and noise in Experiment 1, different head tilts in Experiment 2), we extracted saccade duration and amplitude from the eye tracking data. We averaged these characteristics across trials within each participant, separately for each EMREO condition, and compared the resulting averages between conditions for both experiments across participants.

### 2.8. Quantification of EMREOs

We collected EMREO data for n = 43 participants. However, for the suppression experiment four participants had incomplete datasets (e.g., with one block missing due to corrupt data files or the participant not finishing the experiment), leaving n = 39 complete datasets. For the head tilt experiment all 43 datasets were complete.

For each participant we computed the epoch-averaged EMREO as in previous work [2,3]. The microphone data were aligned to the detected onsets of the largest saccade observed in a 2 s period following the jump of the fixation dot. To avoid artifacts, we only included saccades whose amplitude was within the range 2° to 16°, for which the initial fixation prior to the saccade was stable and for which the entire horizontal and vertical traces of the eye position were contained in an 18° x 18° window around the central fixation point. The resulting data epochs were z-scored to screen for extreme values: epochs for which the maximal z-score of the EMREO signal at any point in time exceeded 4.5 SDs of the overall signal were included in the epoch average. For each ear, we averaged the epochs separately for saccades to the left or right (Experiment 1) or for saccades in each of the four cardinal directions in head coordinates (Experiment 2).

We then checked the SNR of the resulting epoch-averaged EMREO time courses by calculating the ratio of the maximal absolute deflection in a time window of 0 to 50 ms from saccade onset to the standard deviation of the signal in the time window of -150 to 0 ms. Participants with an ear-averaged SNR < 1.5 or an excessive EMREO amplitude (> 20mV) in any of the ears were excluded. Based on these criteria no further participant was excluded for the suppression experiment and four participants for the head-tilt experiment (1 based on the SNR, 3 based on the signal amplitude). As a result, we report data from n = 39 participants for each experiment (though not necessarily the same participants).

### 2.9. Analysis of EMREOs

For the suppression experiment, we asked whether the EMREO time course at any point differs significantly with the presence or absence of the contralateral suppression noise. We performed this comparison separately for the EMREOs induced by saccades to the left and right, given the known difference in EMREO time course for saccades in different directions, and separately for each ear. For this we calculated for each block the epoch-averaged EMREO for all saccades oriented to the left or right, regardless of the starting point of the saccade (center or one of the eccentric positions). This analysis was based on n = 39 participants and an average of 365.8 ± 13.9 (mean, s.e.m.) epochs per participant (reflecting about 77% on average of the recorded data epochs). The statistical analysis was based on a within-participant comparison of conditions using randomization statistics (see below on “Statistical testing”). To summarize the EMREO data, we extracted the average EMREO amplitude around the first main peak (5 to 15 ms) for each participant, ear and saccade direction. To correlate the suppression effect in the EMREO amplitudes with that in the TEOAEs we further averaged these amplitudes for each participant and ear across saccade directions, adjusting for the difference in EMREO sign for saccades to ipsi- and contralateral directions.

The data from the head-tilt experiment were analyzed in two ways. First, we focused on the EMREOs induced by saccades along the four cardinal axes of the head (left, right, up, down in head coordinates) and tested whether these are affected by changes in head tilt. For this we computed the epoch-averaged EMREOs for saccades aligned with the cardinal directions along the head by selecting, for each direction and participant, all saccades whose direction was aligned with the respective cardinal direction in an angular window of ± 10° (in this specific analysis data from 2 participants were excluded because they presented less than 2 trials in one of the directions). This was the case for a subset of the saccades in the remaining 37 participants and this analysis was based on 482.2 ± 13 (mean, s.e.m.) data epochs per participant (reflecting about 67% on average of the recorded data epochs). We then contrasted these EMREOs between the different head tilts. Since the experiment comprised three conditions (no tilt, head-tilt to the left and right) this amounts to a large number of statistical contrasts (2 ears x 4 saccade directions x number of statistical comparisons between conditions x number of time points to be compared). To simplify this, we visualize and describe the data from all three conditions but based the statistical analysis on the comparison of the two head-tilts only. We hence effectively asked, for each ear and saccade direction, whether the EMREO time course differs between tilting the head to the left or right. This statistical analysis was based on a within-participant comparison of conditions using randomization statistics (see below on “Statistical testing”).

In a second analysis we used a regression model to capture the influence of saccade direction and amplitude on the EMREOs, similar to previous work [1,3–5,17,31]. In those studies, it was shown that the EMREO deflection for any saccade can be predicted based on the vertical and horizontal amplitudes and directions of a saccade. Implementing such a regression model allowed us to include all data epochs in the analysis, regardless of the precise saccade direction, and allowed capturing the combined influence of both horizontal and vertical saccade properties and of the tilt manipulation on the EMREO time course in the same analysis. This analysis was based on n = 39 participants and 503.3 ± 14.8 (mean, s.e.m.) data epochs per participant.

We implemented this regression model based on the saccade properties encoded in head-directions, using the following formula, where Sx and Sy reflect the horizontal and vertical saccade amplitude (including its sign) and tilt the experimental block (coded as -1, 0, +1) and t denotes the time point during the EMREO:

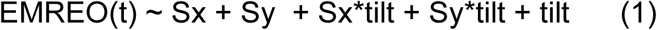

We fit this model separately for each participant, including the data epochs from all three blocks. We then extracted the slopes of each predictor, the overall explained variance (R2) of the model, and the model Akaike Information Criterion (AIC). To quantify the relevance of individual predictors, and to test whether head tilt adds explanatory power to the model, we also computed reduced models that omitted each particular predictor of interest (i.e. Sx, Sy or tilt). From these we derived the unique variance explained by each predictor, defined as the difference in the variance explained by the full and the respective reduced model [1,3–5,17,31]. We also computed the participant-summed AIC values for each model to compare the model-predictive power of each predictor.

### 2.10. Statistical testing

Group-level inference on the EMREO time courses, or the TEOAE spectra, was based on within-participant contrasts and cluster-based permutation tests controlling for multiple comparisons along the dimensions of interest (e.g., frequency for TEOAE spectra or time for EMREOs). First-level contrasts were based on paired t-tests (one-tailed for the TEOAEs; two-tailed for EMREOs). The outcome of these first-level tests was thresholded at p < 0.05 and the test statistic was aggregated across neighboring significant bins (computing the max-sum as cluster-forming statistics and requiring a minimal cluster size of 8 bins)[53]. The resulting clusters in the actual data were compared to potential clusters in 10’000 sets of surrogate data, obtained after randomly flipping the effect-sign for each participant. Clusters with a second-level p < 0.01 were deemed significant. To report a measure of effect size, we report the associated effect size of the underlying within participants t-test (Cohen’s D).

Group-level inference on the EMREO regression analysis was based on the model evidence (AIC values) and the slopes of the individual predictors. To test whether head tilt provides explanatory power on the EMREOs, we contrasted the AIC values between the full model and the model excluding tilt and its interactions with saccades. We converted the difference in AIC values into Akaike model weights, which reflected the conditional probability of each model given the data [54]. To test whether individual factors were significant predictors, we relied on cluster-based permutation tests to test the slope of each factor against zero, correcting for multiple tests along time. Saccade characteristics were compared based on paired t-tests (for amplitude and duration).

When interpreting Cohen’s D effect size, we label effects around D = 0.2 as small, effects around D = 0.5 as medium and D > 0.8 as large. When interpreting Bayes factors we follow the nomenclature of Raftery [55]: we consider BF below 3 as weak evidence, BF between 3 and 20 as positive, BF between 20-150 as strong, and BF above 150 as very strong evidence. Bayes factors larger than 1 indicate evidence that a given predictor adds predictive power, while values smaller than 1 indicate evidence against this hypothesis [56].

## 3. Results

### 3.1. TEOAE suppression

We analyzed TEOAEs and their suppression by a contralateral noise in the frequency domain. Fig 2A shows the group-average TEOAE spectra obtained in silence for each ear. This reveals characteristic power above baseline between about 500Hz and 2 kHz, in line with known properties of TEOAEs [57,58]. Importantly, this TEOAE signal was significantly reduced during presentation of a contralateral noise (Fig 2B). For the left ear, the suppression was significant between 640 and 1280 Hz (n = 39 participants; permutation test, controlling for multiple comparisons along frequency bins; p = 0.0037) with a medium to large effect size (Cohen’s D = 0.59); for the right ear it was significant between 640 and 1472 Hz (p = 0.0005) with a medium effect size (Cohen’s D = 0.52). The insets in Fig 2B show the participant-wise frequency-averaged suppression values (group-level mean ± SD: right ear = 0.94 ± 2.10; left ear = 0.95 ± 2.77), showing that some individuals showed suppression of up to 5 dB.

**Fig 2.**
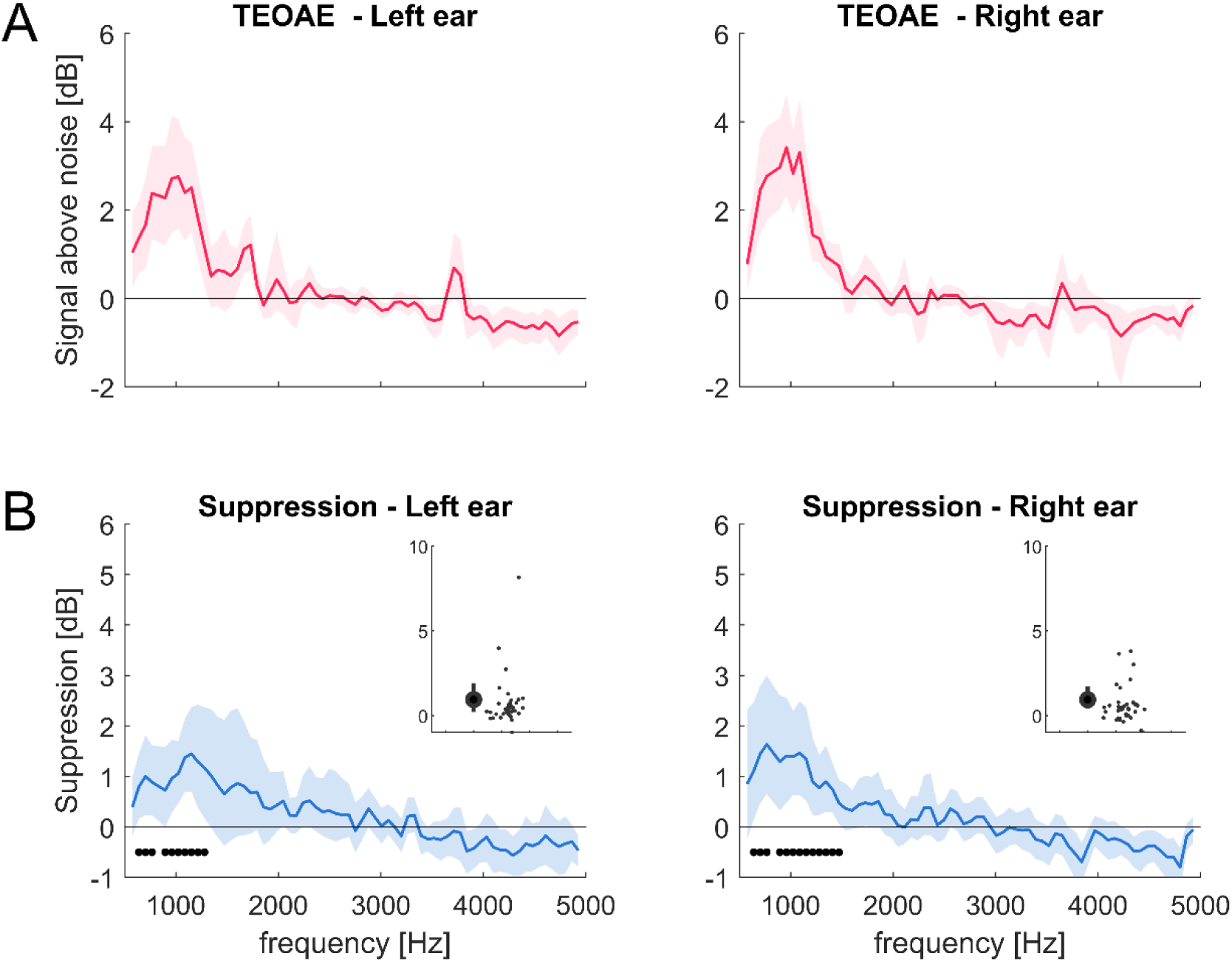
Suppression of TEOAEs. (A) Average TEOAEs spectra above noise for both ears as a function of frequency (pink). Lines indicate group-mean and 95th percentile bootstrap confidence interval (n = 31). (B) Average TEOAE suppression for both ears (blue). Black dots indicate statistically significant suppression (cluster-based permutation test; p < 0.01). Insets indicate the frequency-averaged suppression values, showing the group mean and s.e.m. and the data for individual participants (dots).

### 3.2. No effect of suppression noise on EMREOs

In Experiment 1 we recorded EMREOs for horizontal saccades to the left and right. The EMREO time course followed the previously described pattern, with opposing deflections in the same ear for saccades to opposing directions and lasting about 80 ms (Fig 3A, B; “Silence”). We then tested whether the EMREO time course is altered by the presence of a contralateral noise (Fig 3A, B, “Noise”). A statistical comparison revealed no significant differences for either ear or saccade direction (n = 39 participants; at p < 0.05; corrected for multiple tests along time). To interpret the observed condition-differences, the lower panels in Fig 3A, B show the associated effect sizes (Cohen’s D) for the within participant comparison of the two conditions. These effect sizes were mostly below 0.5, reflecting small effects. The Bayes factors associated with underlying paired t-tests, at the time points of largest effect size, were also small and indecisive (to left and right saccades: left ear BF = 0.18 and 0.45; right ear BF = 0.17 and 0.30). We interpret this as the suppression having at best a small influence on the EMREO time course.

**Fig 3.**
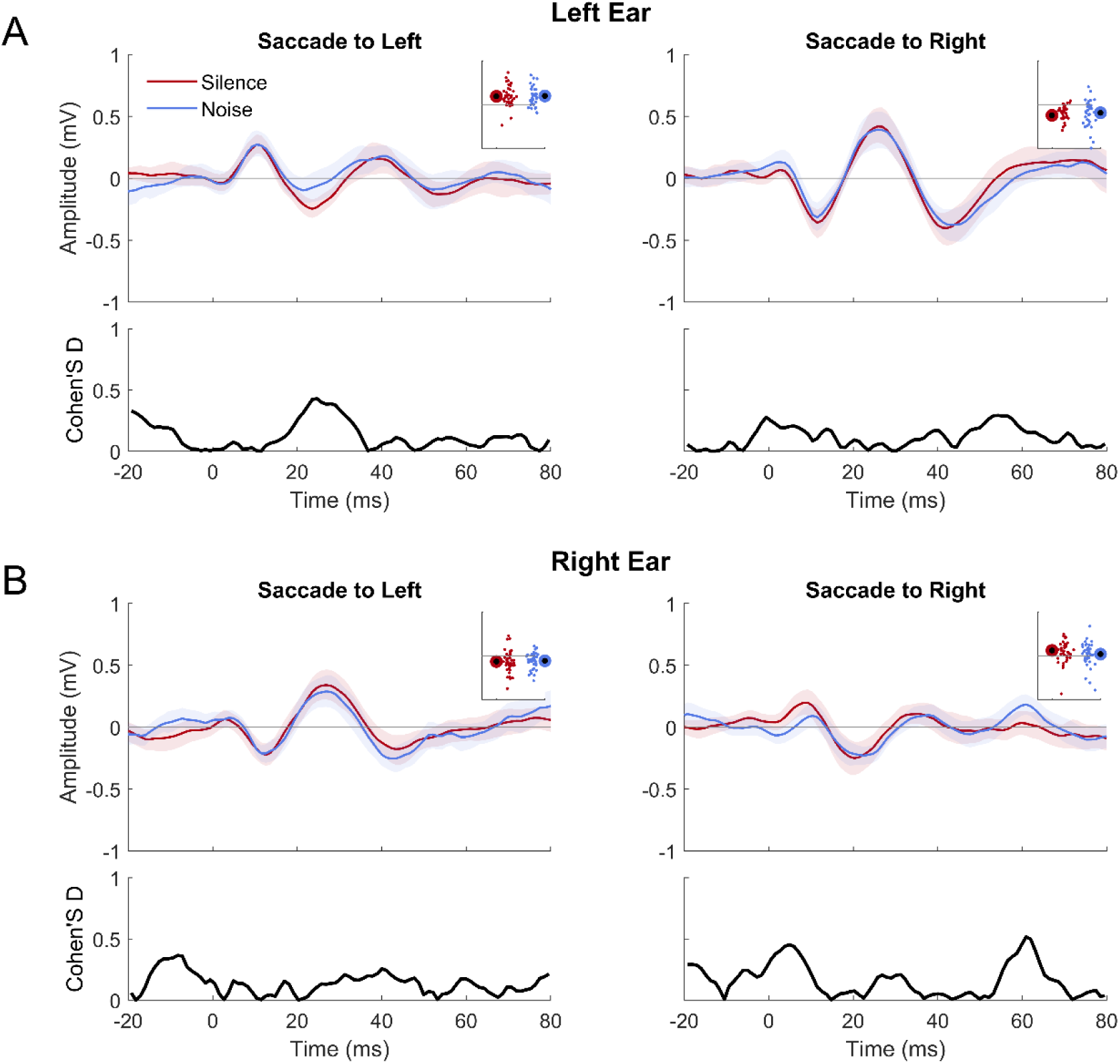
EMREOs for horizontal saccades in silence and with contralateral suppression. The upper panels show the group-average EMREO for the left (A) and right ear (B), split by saccade direction and conditions: saccades in silence (pink) and with contralateral suppression noise (blue). Lines indicate the group-average (n = 39), shaded areas the 95th percentile bootstrap confidence interval. Insets indicate the EMREO amplitudes, showing the group mean and s.e.m. and the data for individual participants (dots). The lower panels show Cohen’s D effect size (black) for the within participant comparison across conditions.

We also tested for an association between the suppression effect seen in the TEOAE and the difference in the EMREO amplitudes. For this we extracted the EMREO amplitude around the initial peak and computed the condition-difference for each ear (mean ± SD: right ear = 0.051±0.17; left ear = 0.002±0.19). Across ears (n = 78) the correlation between the TEOAE and EMREO condition differences were not significant (rho = 0.04, p = 0.69).

### 3.3. EMREOs are not affected by head tilt

For the head tilt task, we analyzed the EMREOS for saccades along the cardinal directions of the head (left, right, up and down) across three conditions (Head Up, Head Left and Head Right). Fig 4 shows the group-level EMREOs for each saccade direction, condition and ear. The time courses for the two horizontal directions reflect the opposing EMREO deflections seen in the suppression paradigm above. The EMREOs for the two vertical directions revealed also consistent modulations but with smaller deflections in line with previous studies [3–5].

**Fig 4.**
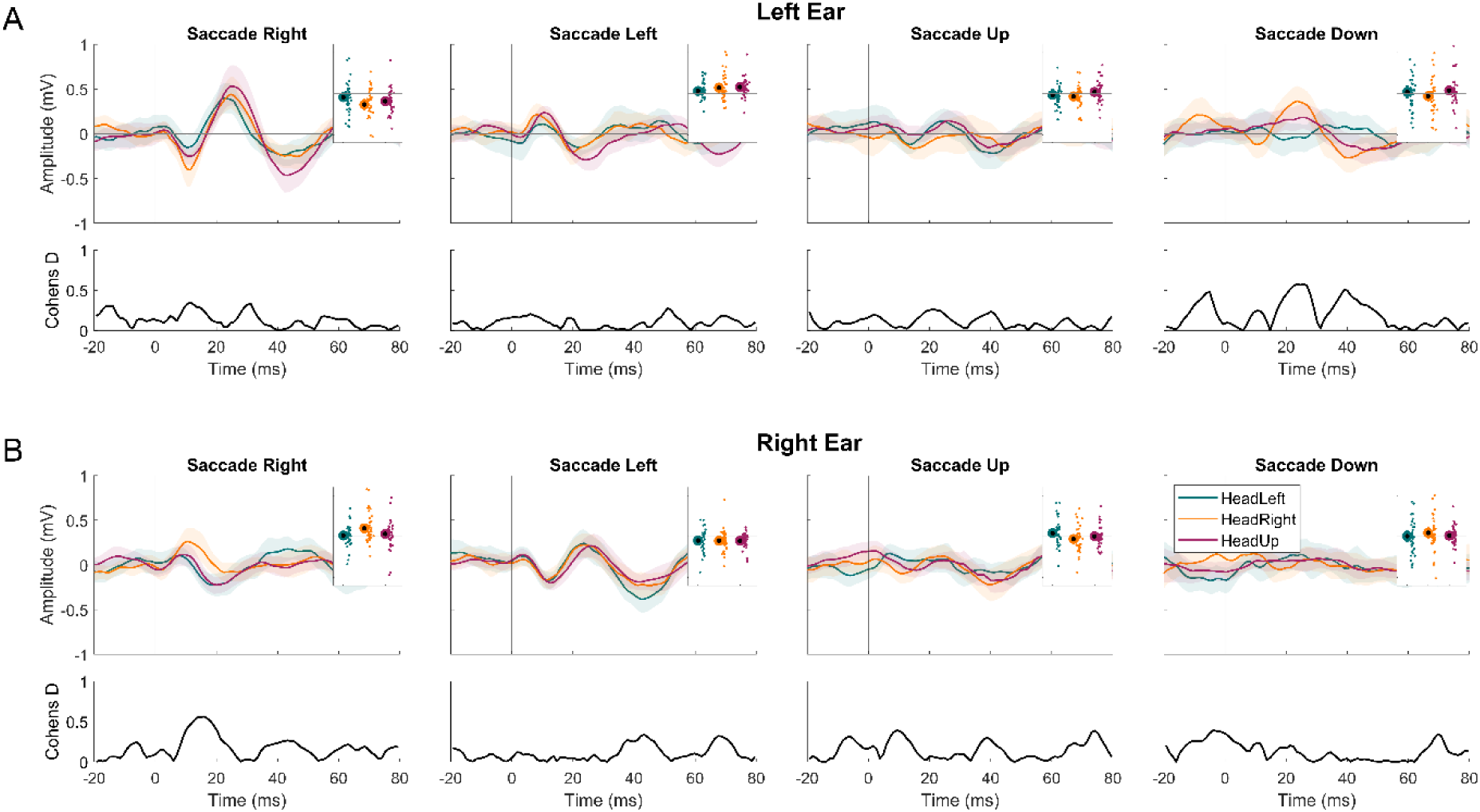
EMREOs in the head-tilt task. The upper panels show the group-average EMREO for the left (A) and right ear (B), split by saccade direction and conditions: head left (green), head right (orange) and head up (red). Lines indicate the group-average (n = 37), shaded areas the 95th percentile bootstrap confidence interval. Insets indicate the EMREO amplitudes, showing the group mean and s.e.m. and the data for individual participants (dots). The lower panels show Cohen’s D effect size (black) for the within participant comparison across conditions.

To test for an influence of head tilt on EMREOs, we contrasted these time courses between the two tilt conditions (tilt left vs. tilt right), separately for each ear and saccade direction. This revealed no significant differences (n = 37 participants; at p < 0.05, corrected for multiple comparisons along time points). Similar as for the suppression paradigm, the associated effect sizes (Cohen’s D) were generally small and reached medium effect size only for a few time points (Fig 4, lower panels). Similar to the suppression experiment, Bayes factors associated with paired t-tests at the largest effect size time points were also small (per direction: left ear BF = 0.61, 0.26, 0.19 and 0.25; right ear BF = 1.58, 0.17, 1.13 and 0.28). To further illustrate this influence of head tilt on EMREOs, Fig 4 insets show the participant-wise EMREOs amplitude for each condition, ear and saccade direction.

In a second analysis, we used regression models to predict the EMREO time course based on the horizontal and vertical saccade amplitude and the head-tilt condition. Fig 5A shows the explained variance of the full regression model. This explained considerable variance, in particular around the major deflections in the EMREO between about 5 and 50 ms as shown previously [3–5,31]. To determine the specific contribution of each model predictor, we computed the unique variance explained by these. This revealed that the horizontal saccade component explained by far the most unique variance, and much less variance was explained by the vertical component (Fig 5A): averaged across ears and the first 50 ms following a saccade, the unique variance explained by the horizontal component was 0.0104±0.0016 (mean ± s.e.m.), that of the vertical component only 0.0015±0.0004.

**Fig 5.**
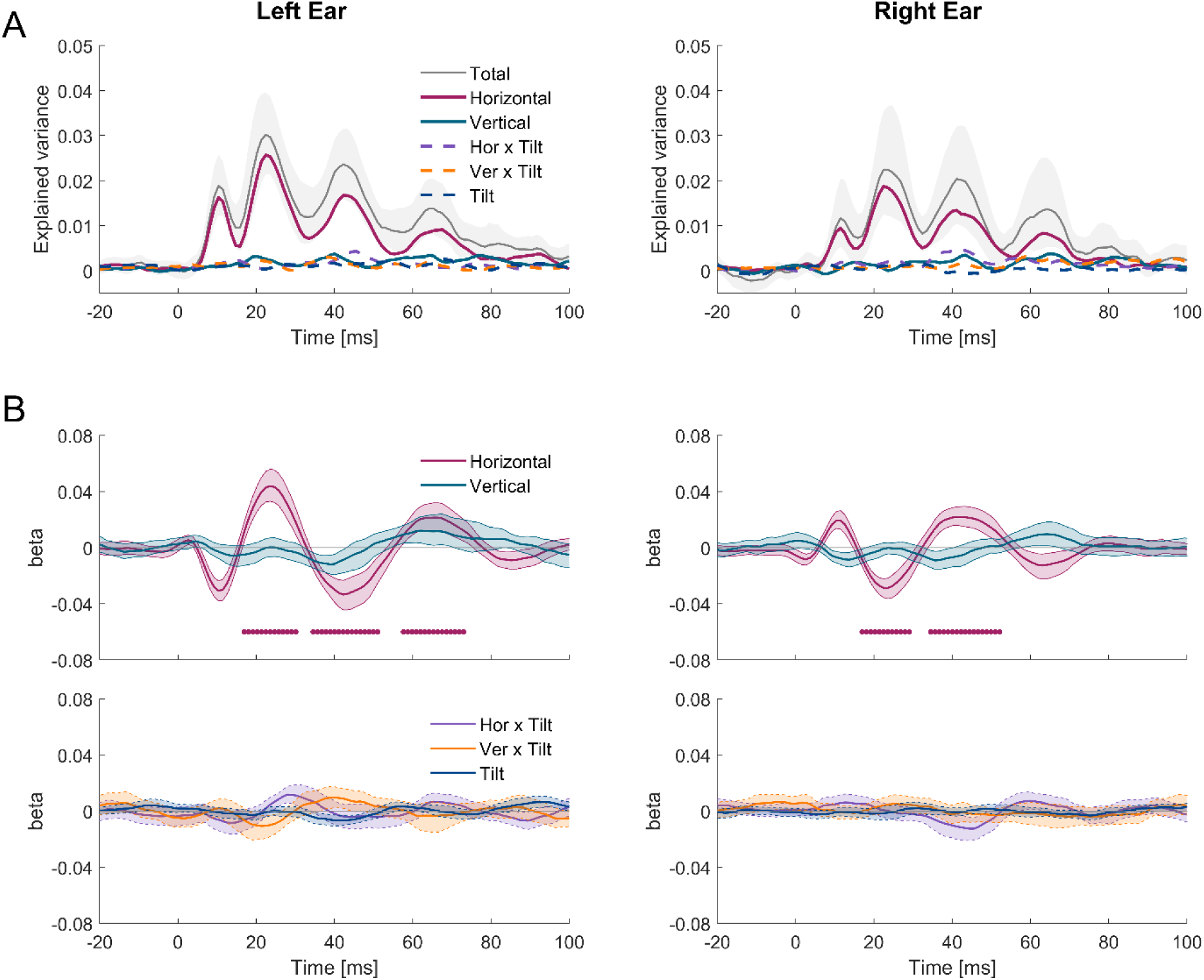
Regression analysis of EMREOs for the Head tilt task. (A) Total variance explained by the full regression model (black) and unique variance explained by individual predictors for ERMEOs in each ear (n = 39). (B) Upper panels show the slopes for the predictors of horizontal (red) and vertical (blue) saccade components. Lower panels show the slopes for tilt condition (green) and its interactions with either the horizontal (yellow) and vertical (purple) saccade components. Black bars indicate statistically significant suppression (cluster-based permutation test; p < 0.01). Lines indicate the group-average time course is and shaded areas the 95th percentile bootstrap confidence interval.

Importantly, also the tilt condition and its interactions with the saccade components explained very little variance (tilt x horizontal: 0.0018±0.0003; tilt x vertical: 0.0011±0.0002; tilt 0.0008±0.0005). For statistical analysis we tested the significance of each model predictor. This revealed multiple clusters of significant time points for the horizontal component (Fig 5B; all p < 0.001) but no significant effects for the vertical component, the tilt or the interactions of tilt and saccade properties (Fig 5B, C; at p < 0.05). Last, we used a model comparison to probe whether knowledge about the tilt condition allowed better predicting the EMREO. This revealed overwhelming evidence that the regression model without tilt explained the EMREO better than a model including tilt (difference in AIC scores: left ear = 217.641, right ear = 216.003; Akaike weight of model without tilt ∼ 1 for both ears). All in all, this provides converging evidence that the EMREOs were not affected by the manipulation of head tilt.

### 3.4. Analysis of saccade properties

Since the EMREOs are shaped by the duration and the amplitude of the inducing saccade, we analyzed the saccade properties for the different experiments and conditions. This was to ensure that there are no differences in saccade parameters that may confound the main comparisons of EMREOs. Fig 6A shows the saccade amplitudes and durations characteristics for the two conditions in the suppression paradigm and Fig 6B, C for the Head tilt experiment.

**Fig 6.**
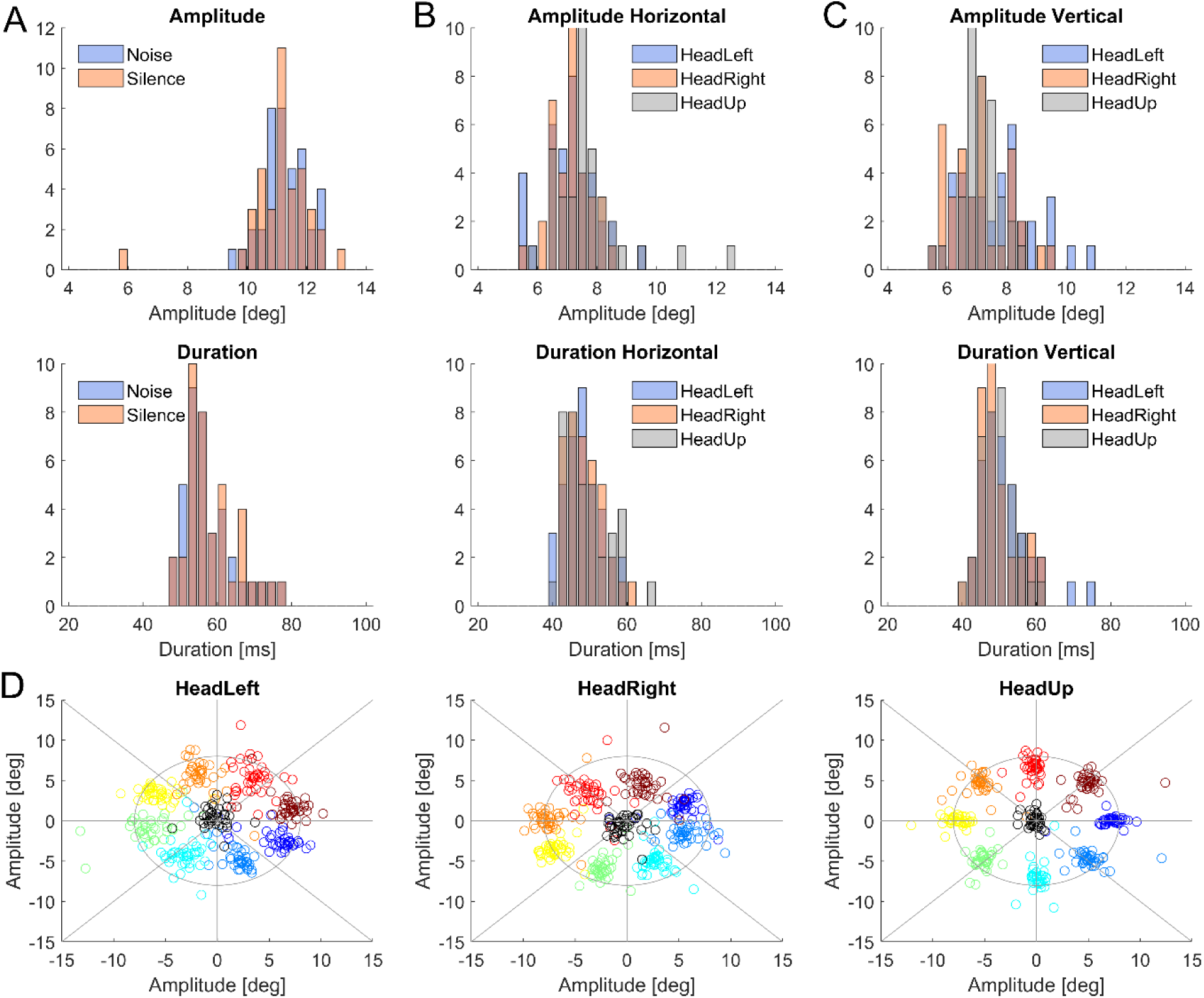
Saccade properties. Participant-wise saccade amplitudes and durations in the different conditions for each paradigm. Panel (A) shows the noise (blue) and silence (orange) conditions for the suppression paradigm. Panels (B) and (C) show the head left (blue), head right (orange) and head up (gray) conditions for in the head-tilt paradigm. Panel (D) shows average saccade end points in head coordinates in the head tilt paradigm (n = 31).

For Experiment 1 (suppression) the statistical tests (ANOVA) revealed no significant effects of condition on either the saccade amplitude (n = 39 participants; F(76) = 0.7219, p = 0.398) or duration (F = 0.145, p = 0.704). For Experiment 2 (head tilt) this revealed a borderline significant effect of saccade direction (n = 39 participants; F(145) = 4.03, p = 0.046) and an effect of condition on saccade amplitude (F = 4.79, p = 0.03). Saccade durations did differ between saccade directions (F = 11.52, p = 0.0008) but not between conditions (F = 1.032, p = 0.31). Hence, overall saccades did not differ between the two tilt conditions. Fig 6D shows the average saccade landing points in the head tilt experiment for each participant and condition. This reveals a rotation of the end points with the orientation of the head, as expected given the experimental manipulation. However, as observed in a previous study, the resulting distribution of end points does not completely follow the head rotation and reveals larger variability and a compression along the vertical direction [59].

## 4. Discussion

While EMREOs have been reported in a number of studies, their properties and the underlying neural pathways giving rise to this phenomenon remain unclear. We here tested for an effect of two experimental manipulations that are known to either influence cochlear activity or which are likely to alter body- and oculomotor-related feedback signals to the ear, and hence could possibly influence the EMREO phenomenon. However, neither the presentation of a contralateral suppression noise, which reduced TEOAEs, nor tilting the head by a significant amount, induced clear and significant changes in the EMREO. All in all, this suggests that the EMREO signal reflects a robust and temporally stable signal.

### 4.1. Known properties of EMREOs

The first systematic description of the EMREO phenomenon was made based on recording in humans and monkeys in silence in 2018 [1]. Since then, several studies have corroborated this phenomenon and have described a number of properties of the EMREO signal recorded in the ear canal that we summarize in the following.

EMREOs emerge for saccades in all directions (i.e., for the horizontal, vertical and diagonal dimensions) [1,4,31], but the EMREO signal is more reliable and more strongly modulated for horizontal compared to vertical saccades [4,31]. The amplitude and the time course of vertical EMREOs shows higher inter-individual variability, and shows no reliable pattern of phase opposition compared to horizontal saccades [4]. Our data corroborate this observation as they reveal a stronger EMREO amplitude and a more pronounced temporal structure for horizontal compared to vertical saccades (c.f. Fig 5). The interpretation of this difference in EMREO properties for horizontal and vertical saccades remains unclear, and requires further investigation.

The EMREO for horizontal saccades is more time-locked to saccade onset than to offset [1,4,31] and generally comparable in the left and right ears [1,2]. However, within each ear, the EMREO time course differs for saccades to targets ipsi- and contralateral to the ear under study, and the signal starts with deflections of opposing signs for the two saccade directions. Although this results in a pattern of apparent phase-opposition between saccade directions, the eardrums in the two ears don’t move in strict antiphase for a given saccade: accounting for the difference in EMREO sign between saccades to ipsi- and contralateral targets still leaves a significant difference in the signal time course between saccade directions [2]. Hence, for any given horizontal saccade, the underlying movements of the tympanic membranes in the left and right ears differ, resulting in a net-difference between the state of the two eardrums over time.

Still, across all saccade directions the EMREO signal can be modelled based on knowledge of the saccade target [4,5,31]. Our regression analysis directly corroborates this, but also suggests that the horizontal saccade amplitude is a more reliable predictor of the EMREO amplitude compared to the vertical saccade amplitude. The observation of this relation between saccade properties and EMREO led to the suggestion that the EMREO signal may reflect some form of spatial coordinate transform between eye and ear centered maps, possibly playing a role in coordinating the encoding of multisensory signals [1,4,5]. However, the accuracy of such a coordinate transformation remains unclear, given that the scaling of EMREO amplitude with saccade eccentricity does not seem to be linear [4].

While many studies recorded EMREOs during visually-guided saccades, EMREOs also emerge during free-viewing [31], auditory-guided saccades [2], in darkness [17,60] and with the eyes closed [60]. The EMREO time course appears very similar during free-viewing and auditory induced saccades compared to visually-guided saccades [2,31], suggesting that the EMREO phenomenon is tied to the execution of a saccade and not the specific sensory signal or goal driving the motor command. This is also supported by a study reporting that a visual cuing signal prior to a saccade was successful in increasing perceptual sensitivity, but not changing the EMREOs [3], and by studies showing that the EMREO signal is not affected by concurrent acoustic stimuli that are not serving as saccade targets [1,3].

Given that the eardrums in the left and right ears move largely in opposite directions but not in strict antiphase, studies have asked whether this relative movement of the two eardrums has a consequence for auditory perception. One study tested this for a task requiring participants to detect a faint click presented before, during or after the saccade. However, this found no impact of the sound-to-EMREO relation on detection performance [3]. A second study tested participants’ ability to judge the relative horizontal position of two tones, one of which was presented at different time points during the EMREO. However, also in this task, which required participants to exploit spatial acoustic information, there was no significant influence of the timing of the sounds relative to the EMREO time course on behavioral performance [45]. Hence while the emergence of EMREOs is a very robust phenomenon, its relation to hearing or perception in general remains elusive.

Similar to the potential impact of EMREOs on perception, also their origin remains unclear. Prominent candidates include the outer hair cells and the middle ear muscles. However, their specific contribution to the EMREO signal remains to be elucidated. The motivation of the present study was that interfering with their normal function, or otherwise stimulating the pathways activating these, should also modulate the EMREO signal. However, as discussed in the following this was not the case.

### 4.2. EMREOs and outer hair cells

In the first experiment, we tested whether inducing the medial olivocochlear reflex (MOCR) would affect the EMREO signal. During the MOCR, the contralateral presentation of a noise reduces OHC activity in the opposite ear and this reduces the amplitude of transient otoacoustic emissions (TEOAEs). In our data, this suppression was prominent between about 640 and 1472 Hz in the frequency domain. This is consistent with previous studies showing that the MOCR is most prominent at the low-frequency end of the human hearing spectrum [57,58]. The average TEOAE suppression was between 0.94 dB (right ear) and 0.95 dB (left ear), values that are comparable to reports in the literature [15,61–63].

The suppression of OAEs by contralateral noise seems robust to changes in task context, such as during performance of an additional visual task [29,64]. However, large changes in gaze direction were reported to modulate the degree of contralateral suppression of TEOAEs [10]. This points to an influence of oculomotor related signals on the medial olivocochlear system mediating the suppression. Thus, if the EMREOs were generated by the same pathways giving rise to the suppression of OAEs by contralateral noise, one could speculate that the same suppression should also affect the eye movement-related signals emanating from the ear, such as EMREOs. However, our data do not support this.

In fact, noise that consistently reduced TEOAEs did not significantly affect the EMREO time course. One interpretation is that this directly speaks against the OHC as the central effector producing the EMREO phenomenon. This would be consistent with the observation that recordable acoustic signals in the ear generated by OHCs (e.g., otoacoustic emissions) are much smaller in amplitude than EMREOs [1]. In addition, EMREOs are a signal that has most temporal structure around 20-40Hz, whereas TEOAEs are typically observed at a much higher time scale of above 500 Hz, possibly because both signals are reflecting mechanistically distinct processes in the ear. Of course, the difference in time scales does not by itself preclude a common underlying generator, as a neural or mechanical process giving rise to a fast signal could also be modulated at a slower time scale. However, at present the direct evidence for an involvement of OHCs in the EMREO phenomenon remains weak.

In this regard one should note that individual thresholds for the MOCR vary considerably across individuals and their measurement is always associated with some uncertainty [16,25,61]. Hence, the strength of the actually induced neural and functional suppression may have varied somewhat across individuals, adding uncertainty to this experimental manipulation. Despite this individual variability and uncertainty, one would expect that if the EMREO phenomenon is mainly driven by OHCs, any change in EMREO amplitude is associated with the reduction in TEOAEs. However, in our data this was not the case. This uncertainty is also present in other measurable processes used as biomarkers for assessing hearing function, such as OAEs themselves. We therefore believe that, although this disparity should be taken into account, it should not undermine the relevance of our results.

In addition, dissociating the influences of OHCs and MEMs is inherently difficult. Some studies suggest that also the product of OAEs as measured in the ear canal may reflect contributions from both the cochlea and the middle ear [65,66]. Data from patients shows that the stapedius reflex does not interfere with the contralateral suppression of TEOAEs, but cannot rule out contributions of the tensor tympani to this [27]. Hence while the data from the suppression paradigm support the notion that the EMREO signal is shaped to a strong part by the middle ear and not the OHCs, dissociating the contribution of different elements in the ear remains a challenge.

### 4.3. EMREOs and head-tilts

The oculomotor, auditory and vestibular systems are intricately connected and information about our head position profoundly contributes to the execution of saccades [41,67]. This includes the vestibulo-ocular reflex (VOR), which activates the eye muscles contingent on the head position [38,42] and which can induce rotations of the eyes that compensate opposite movements of the head [68]. Vestibular and postural information related to head, neck and body movements contributes to stability of gaze during movements [38,40,59]. This information also reaches the auditory pathways [7,69], where it presumably stimulates the MEMs motor neurons, shaping the acoustic reflex [7,36] and possibly modulating the EMREO. However, whether vestibular and other postural signals related to change in head tilt modulate the EMREO has not been studied before.

Our data support that EMREOs are robust to such an influence of vestibular and proprioceptive signals despite vestibular-ocular reflexes likely operating during the head tilt. The absence of a head tilt effect on the EMREO waveform indicates that the encoding of saccades parametric information is fixed to the head reference, rather than to the body or the visual scene. This interpretation does not support the idea that EMREOs would provide retinocentric spatial information for an adjustment of the computation of sound position. While this finding does constrain our understanding of how OHCs or MEMs contribute to the EMREO, it raises questions about how the underlying processes could help in coordinating visual and auditory signals during natural free viewing.

## 5. Conclusion

The processes underlying the EMREO signal have been implicated in contributing to the coordination of spatial information between the visual and auditory senses [5]. However, such a role seems only partially plausible given that the EMREO signal is shaped mostly by the direction of a saccade (ipsi- or contralateral to the ear under study) and not the precise amplitude [4], given that vertical EMREOs seem less reliable than horizontal ones and given that the position of the head is not factored in. A recent study now offers an alternative idea, speculating that the processes underlying the EMREO may serve more the temporal coordination of multisensory signals than the spatial coordination of these [70]. In fact, it has long been known that both eye movements and visual stimuli shape rhythmic activity along the auditory pathways and temporally adjust the excitability of thalamic and cortical auditory regions [71–73]. One physiological mechanism underlying this form of temporal multisensory interaction is a phase reset of ongoing rhythmic activity, which supposedly serves to coordinate the temporal encoding of sensory signals between different senses [74,75]. Intriguingly, rhythmic activity not only prevails in auditory cortical circuits but has recently been revealed in the ear itself [76,77] and exhibits similar attention-related multisensory effects as seen in cortical activity [78–80]. Hence one possibility is that the EMREO signal is reflecting neural processes that serve the temporal coordination of the encoding of multisensory signals. However, as other speculations about the EMREO phenomenon, this requires careful future work.

## Acknowledgements

We would like to thank Alexander Wecker and the Mechatronics Workshop of Bielefeld University for their support in designing and building the head-tilting platform.

## Author contributions

NSS and CK Conceived and Designed research. NSS Performed experiments. NSS and CK Analyzed data, Interpreted results of experiments, Prepared figures, Drafted and Edited the manuscript. CK Revised and Approved final version of the manuscript

